# Statistics of the number of equilibria in random social dilemma evolutionary games with mutation

**DOI:** 10.1101/2021.07.13.452172

**Authors:** Manh Hong Duong, The Anh Han

## Abstract

In this paper, we study analytically the statistics of the number of equilibria in pairwise social dilemma evolutionary games with mutation where a game’s payoff entries are random variables. Using the replicator-mutator equations, we provide explicit formulas for the probability distributions of the number of equilibria as well as other statistical quantities. This analysis is highly relevant assuming that one might know the nature of a social dilemma game at hand (e.g., cooperation vs coordination vs anti-coordination), but measuring the exact values of its payoff entries is difficult. Our delicate analysis shows clearly the influence of the mutation probability on these probability distributions, providing insights into how varying this important factor impacts the overall behavioural or biological diversity of the underlying evolutionary systems.

## 1 Introduction

### 1.1 The replicator-mutator equation

The replicator-mutator equation is a set of differential equations describing the evolution of frequencies of different strategies in a population that takes into account both selection and mutation mechanisms. It has been employed in the study of, among other applications, population genetics [11], autocatalytic reaction networks [29], language evolution [21], the evolution of cooperation [14, 20] and dynamics of behavior in social networks [22].

Suppose that in an infinite population there are *n* types/strategies *S*_1_, …, *S_n_* whose frequencies are, respectively, *x*_1_, …, *x_n_*. These types undergo selection; that is, the reproduction rate of each type, *S_i_*, is determined by its fitness or average payoff, *f_i_*, which is obtained from interacting with other individuals in the population. The interaction of the individuals in the population is carried out within randomly selected groups of *d* participants (for some integer *d*). That is, they play and obtain their payoffs from a *d*-player game, defined by a payoff matrix. We consider here symmetric games where the payoffs do not depend on the ordering of the players in a group. Mutation is included by adding the possibility that individuals spontaneously change from one strategy to another, which is modeled via a mutation matrix, *Q* = (*q_ji_*), *j, i* ∈ {1, …, *n*}. The entry *q_ji_* denotes the probability that a player of type *S_j_* changes its type or strategy to *S_i_*. The mutation matrix *Q* is a row-stochastic matrix, i.e.,

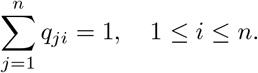

The replicator-mutator is then given by, see e.g. [17, 15, 16, 24]

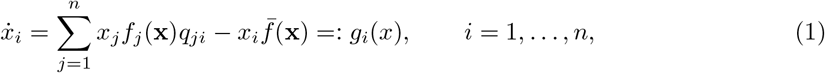

where **x** = (*x*_1_, *x*_2_, …, *x_n_*) and 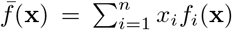 denotes the average fitness of the whole population. The replicator dynamics is a special instance of (1) when the mutation matrix is the identity matrix.

### 1.2 The replicator-mutator equation for two-player two-strategy games

In particular, for two-player two-strategy games the replicator-mutator equation is

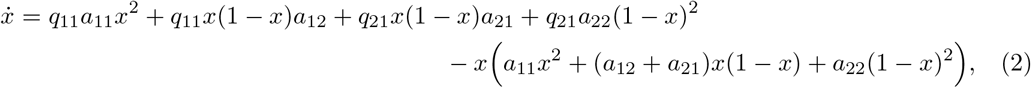

where *x* is the frequency of the first strategy and 1 − *x* is the frequency of the second one. Using the identities *q*_11_ = *q*_22_ = 1 − *q*, *q*_12_ = *q*_21_ = *q*, Equation (2) becomes

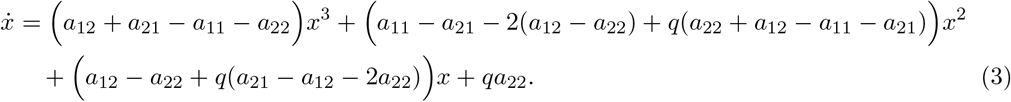

#### Two-player social dilemma games

In this paper, we focus on two-player (i.e. pairwise) social dilemma games. We adopt the following parameterized payoff matrix to study the full space of two-player social dilemma games where the first strategy is cooperator and second is defector [26, 32, 30], *a*_11_ = 1; *a*_22_ = 0; 0 ≤ *a*_21_ = *T* ≤ 2 and −1 ≤ *a*_12_ = *S* ≤ 1, that covers the following games

i. the Prisoner’s Dilemma (PD): 2 ≥ *T* > 1 > 0 > *S* ≥ −1,
ii. the Snow-Drift (SD) game: 2 ≥ *T* > 1 > *S* > 0,
iii. the Stag Hunt (SH) game: 1 > *T* > 0 > *S* ≥ −1,
iv. the Harmony (H) game: 1 > *T* ≥ 0, 1 ≥ *S* > 0.

In this paper, we are interested in random social dilemma games where *T* and *S* are uniform random variables in the corresponding intervals, namely

- In PD games: *T* ~ *U* (1, 2), *S* ~ *U* (−1, 0),
- In SD games: *T* ~ *U* (1, 2), *S* ~ *U* (0, 1),
- In SH games: *T* ~ *U* (0, 1), *S* ~ *U* (−1, 0),
- In H games: *T* ~ *U* (0, 1), *S* ~ *U* (0, 1).

Random evolutionary games, in which the payoff entries are random variables, have been employed extensively to model social and biological systems in which very limited information is available, or where the environment changes so rapidly and frequently that one cannot describe the payoffs of their inhabitants’ interactions [4, 3, 19, 12, 5, 6, 10, 8]. Equilibrium points of such evolutionary system are the compositions of strategy frequencies where all the strategies have the same average fitness. Biologically, they predict the co-existence of different types in a population and the maintenance of polymorphism.

In this paper, we are interested in computing the probability distributions of the number of equilibria, which is a random variable, in the above random social dilemmas. Answering this question is of great importance in the context of social dilemmas since one might know the nature of the game, i.e. the payoff entries ranking in the game, but it might be difficult to predict or measure the exact values of these entries. When mutation is absent (*q* = 0), the answer is trivial because there is always a fixed number of equilibria depending on the nature of the social dilemmas [20, 27, 7]. As shown in our analysis below, this is however not the case any longer when mutation is non-negligible, and this number highly depends on the nature of the social dilemma too.

The following result [7] provides partial information about these probability distributions.

##### Theorem 1.1.

*(DuongHanDGA2020) Suppose that S and T are uniformly distributed in the corresponding intervals as above. Then*

- 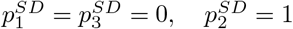
- 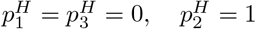
- 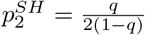
- 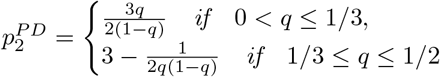

According to the above theorem, SD games and H games are simple and complete. However, the probabilities of having 3 equilibria (or alternatively 1 equilibrium) in SH and PD games are left open in [7]. The key challenge is that the conditions for these games to have 3 equilibria (or alternatively 1 equilibrium) are much more complicated than those of 2 equilibria. The aim of this paper is to complete the above theorem, providing explicit formulas for these probabilities. As such, it will also allow us to derive other statistical quantities (such as average and variance), which are important to understand the overall distribution and complexity of equilibrium points in pairwise social dilemmas (with mutation). To this goal, we employ suitable changes of variables, which transform the problem of computing the probabilities to calculating areas, and perform delicate analysis.

### 1.3 Main results

The main result of this paper is the following.

#### Theorem 1.2.

*The probability that SH and PD games have* 3 *equilibria is given by, respectively*

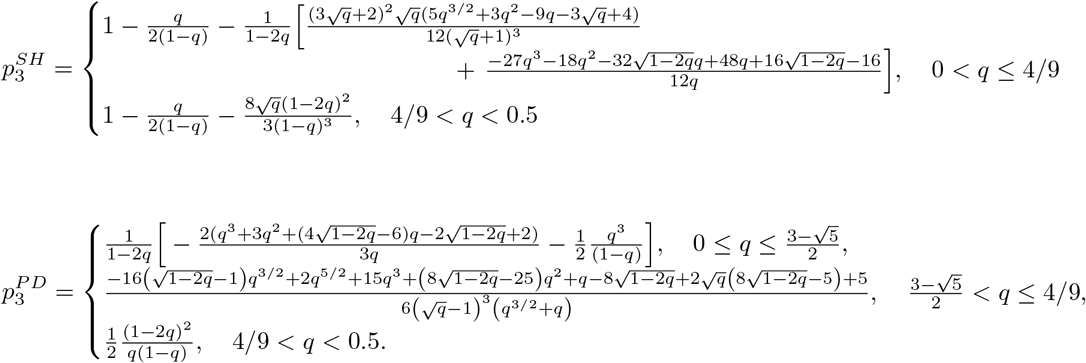

The above theorem combines Theorem 2.3 (for SH-games) and Theorem 2.6 (for PD games), see Section 2. Theorems 1.1 and 1.2 provides explicitly the probability distributions of the number of equilibria for all the above-mentioned pairwise social dilemmas. In SH-games and PD-games, these distributions are much more complicated and significantly depend on the mutation strength. We summarize these results in the following summary box.

#### Box 1

##### Probability of having *k* equilibria in a pairwise social dilemma (*p_k_*)

- **Snow Drift (SD)**

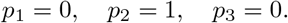
- **Harmony game (H)**

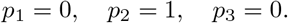
- **Stag-Hunt game (SH)**

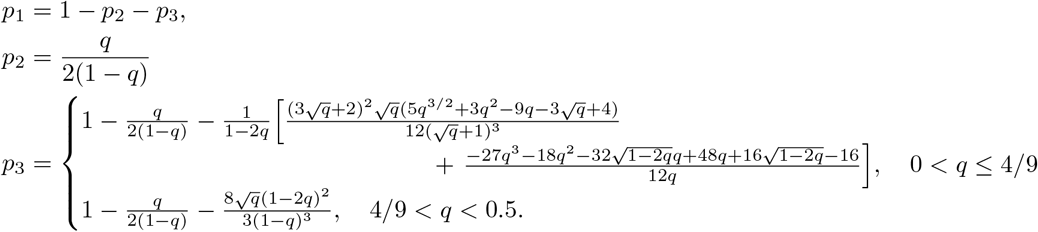
- **Prisoner’s Dilemma (PD)**

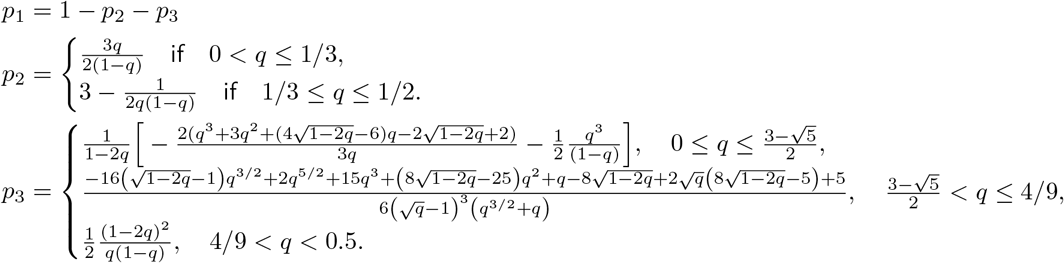

As a consequence, we can now derive other statistical quantities such as the mean value, ENoE, and the variance, VarNoE of the number of equilibria using the following formulas

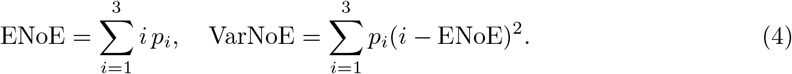

We depict these quantities in Figure 1 below. Our delicate analysis clearly shows the influence of the mutation on the probability distributions, thus on the complexity and bio-diversity of the underlying evolutionary systems. We believe that our analysis may be used as exemplary material for teaching foundational courses in evolutionary game theory, computational/quantitative biology and applied probability.

**Figure 1:**
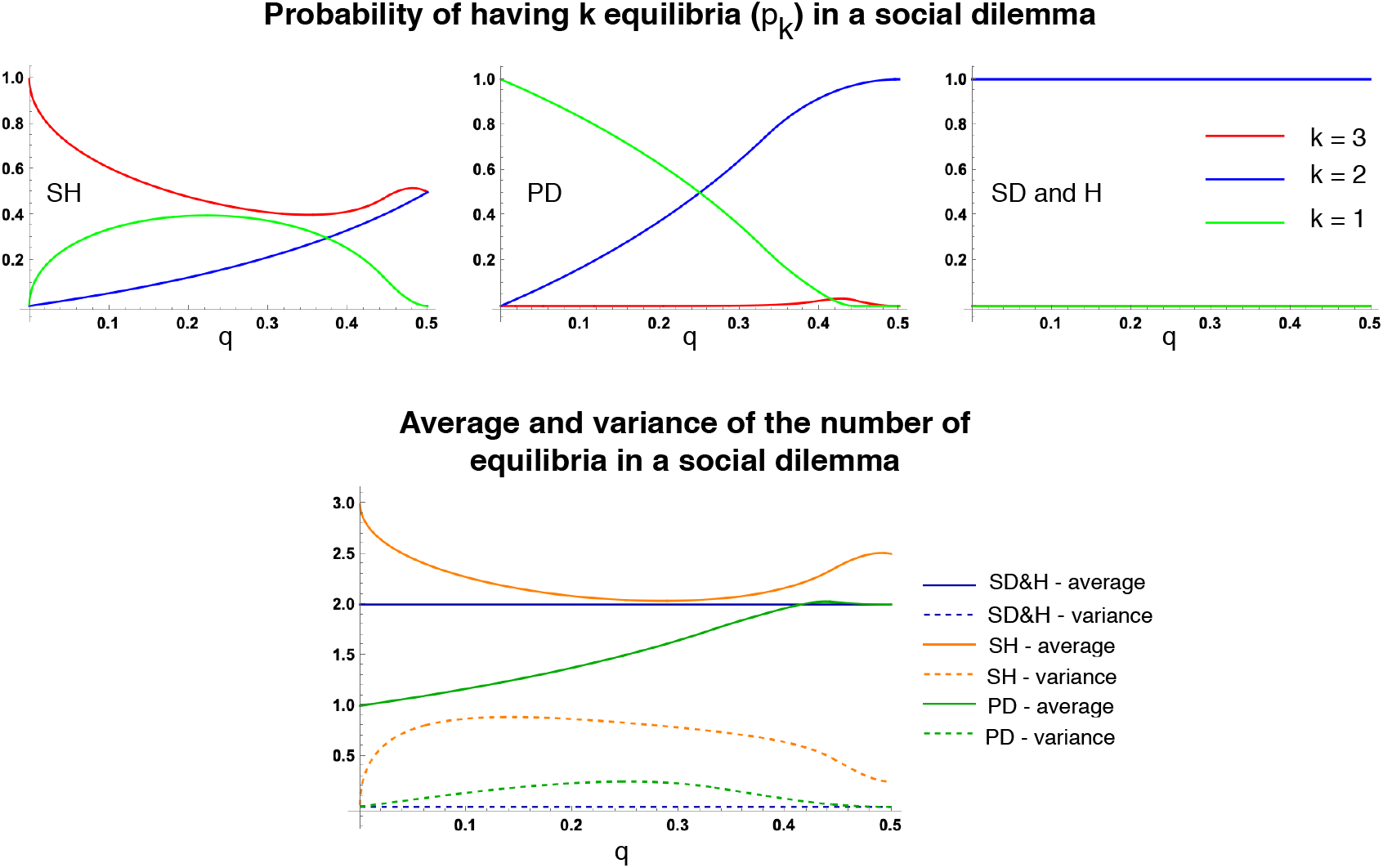
Statistics of the number of equilibria in random social dilemmas as a function of the mutation probability *q*: probability of having *k* (*k* = 1, 2, 3) equilibria (**top row**), average and variance (**bottom row**). We look at all the four pairwise games, from left to right: SH, PD, SH and H. The probability of having the maximal possible number of equilibria, i.e. *p*_3_, is highest for SH. It is very small for PD and always equals 0 for SD and H games. The probability of having two equilibria, *p*_2_, is highest for SD and H. As a result, SH has the highest average number of equilibria across all games, for all 0 < *q* < 0.5 (it is minimum when *q* = 0.285). For most *q*, PD has the lowest number of equilibria (it is maximum when *q* = 0.438). All the figures are generated using analytical formulas derived in Box 1 (for *p_k_*) and (4) for the average and variance. These analytical results are also in accordance with numerical simulation results provided in [7], obtained through samplings of the random payoff entries *T* and *S*.

The rest of the paper is organized as follows. In Section 2, after recalling some preliminary details, we present the proof of the main theorem 1.2, which we split into Theorem 2.3 for SH games in subsection 2.3 and Theorem 2.6 for PD games in subsection 2.4. Finally, in Section 3 we provide further discussion.

## 2 Probability of having three equilibria in SH and PD games

### 2.1 Joint probability density via change of variable

The following lemma is a well-known result to compute the probability density of random variables using change of variables. We state here for two random variables that are directly applicable to our analysis, but the result is true in higher dimensional spaces.

#### Lemma 2.1.

*(joint probability density via change of variable, [9, Section 3.3]) Suppose (X_1_, X_2_) has joind density f (x_1_, x_2_). Let (Y_1_, Y_2_) be defined by Y_1_ = u_1_(X_1_, X_2_) and Y_2_ = u_2_(X_1_, X_2_). Suppose that the map (X_1_, X_2_) → (Y_1_, Y_2_) is invertible with X_1_ = v_1_(Y_1_, Y_2_) and X_2_ = v_2_(Y_1_, Y_2_). Then the joint probability distribution of Y_1_ and Y_2_ is*

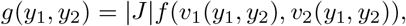

*where the Jacobian J is given by*

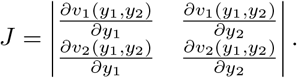

### 2.2 Equilibria in Social dilemmas

By simplifying the right hand side of (3), equilibria of a social dilemma game are roots in the interval [0, 1] of the following cubic equation

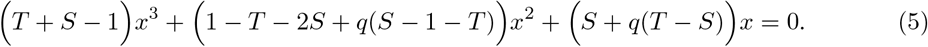

It follows that *x* = 0 is always an equilibrium. If *q* = 0, (5) reduces to

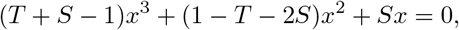

which has solutions

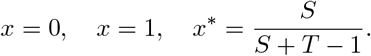

Note that for SH-games and SH-games *x*\* ∈ (0, 1), thus it is always an equilibrium. On the other hand, for PD-games and H-games, *x*\* ∉ (0, 1), thus it is not an equilibrium

If 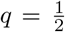 then the above equation has two solutions 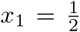 and 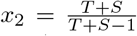. In PD, SD and H games, *x*_2_ ∉ (0, 1), thus they have two equilibria *x*_0_ = 0 and 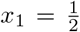. In the SH game: if *T* | *S* < 0 then the game has three equilibria *x*_0_ = 0, 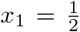 and 0 < *x*_2_ < 1; if *T* | *S* ≥ 0 then the game has only two equilibria *x*_0_ = 0, 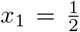.

Now we consider the case 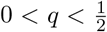. For non-zero equilibrium points we solve the following quadratic equation

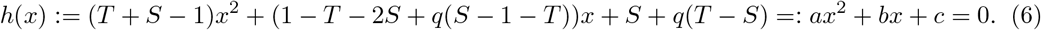

Set 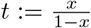, then we have

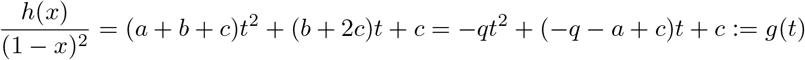

Therefore, a social dilemma has three equilibria iff *h* has two distinct roots in (0, 1), which is equivalent to *g* having two distinct positive roots, or the following conditions must hold

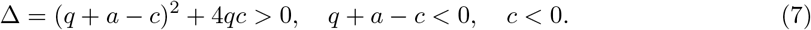

Let 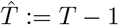 and

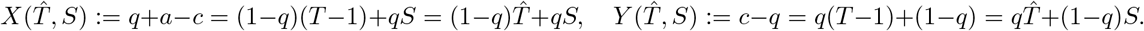

The inverse transformation 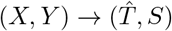 is given by

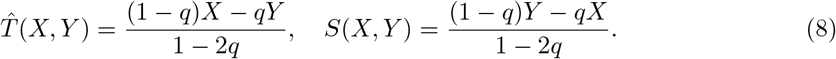

Condition (7) is given by

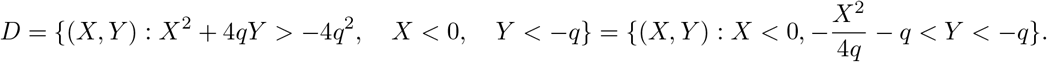

The domain *D* is illustrated in Figure 2. We apply Lemma 2.1 to find the joint distribution of *X* and *Y*. We compute the Jacobian of the transform 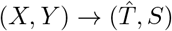, which is given by

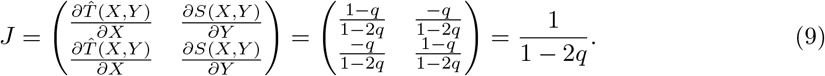

**Figure 2:**
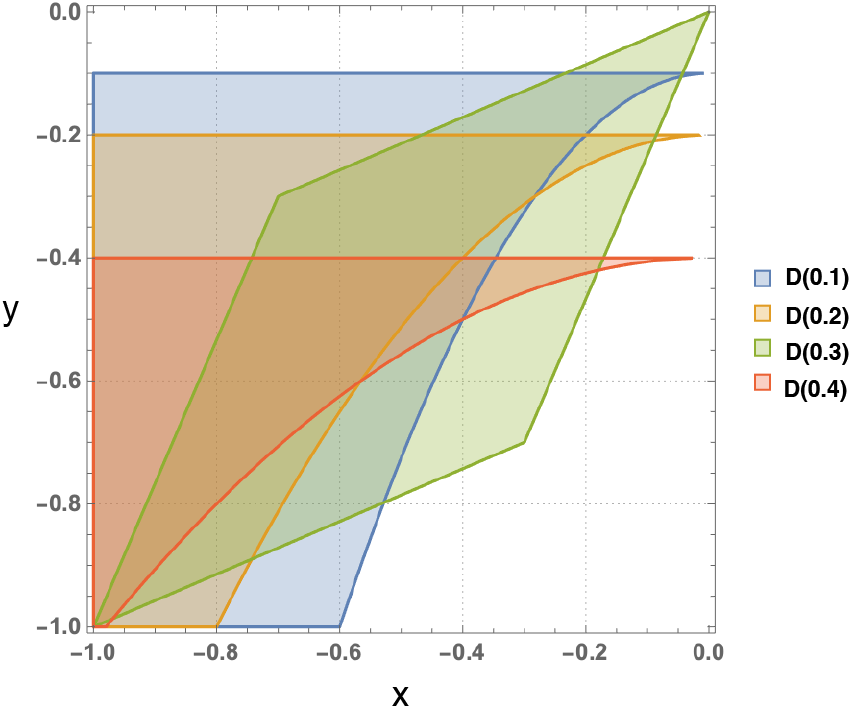
Region *D* are shown for different values of *q*, namely, *q* = 0.1, 0.2, 0.3 and 0.4.

Hence if 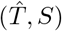 has a probability density *f* (*t, s*) then (*X, Y*) has a probability density

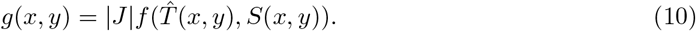

We now apply this approach to SH and PD games.

### 2.3 The Stag Hunt (SH)

#### Proposition 2.2.

*The probability that SH games have* 3 *equilibria is given by*

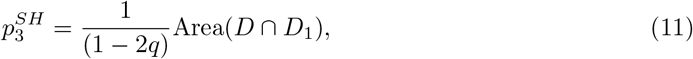

*where D is the subset of* ℝ^2^ *determined by*

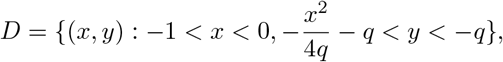

*and D_1_ is the quadrilateral ABOC with vertices*

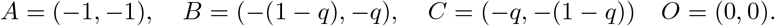

The domain *D*_1_ and the intersection *D* ∩ *D*_1_ is illustrated in Figure 3.

*Proof.* In the SH game: 1 > *T* > 0 > *S* > − 1, *T* ~ *U* (0, 1), *S* ~ *U* (−1, 0). Let 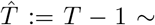 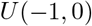. The joint distribution of 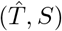 is

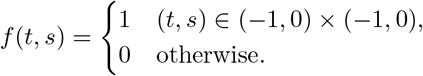

**Figure 3:**
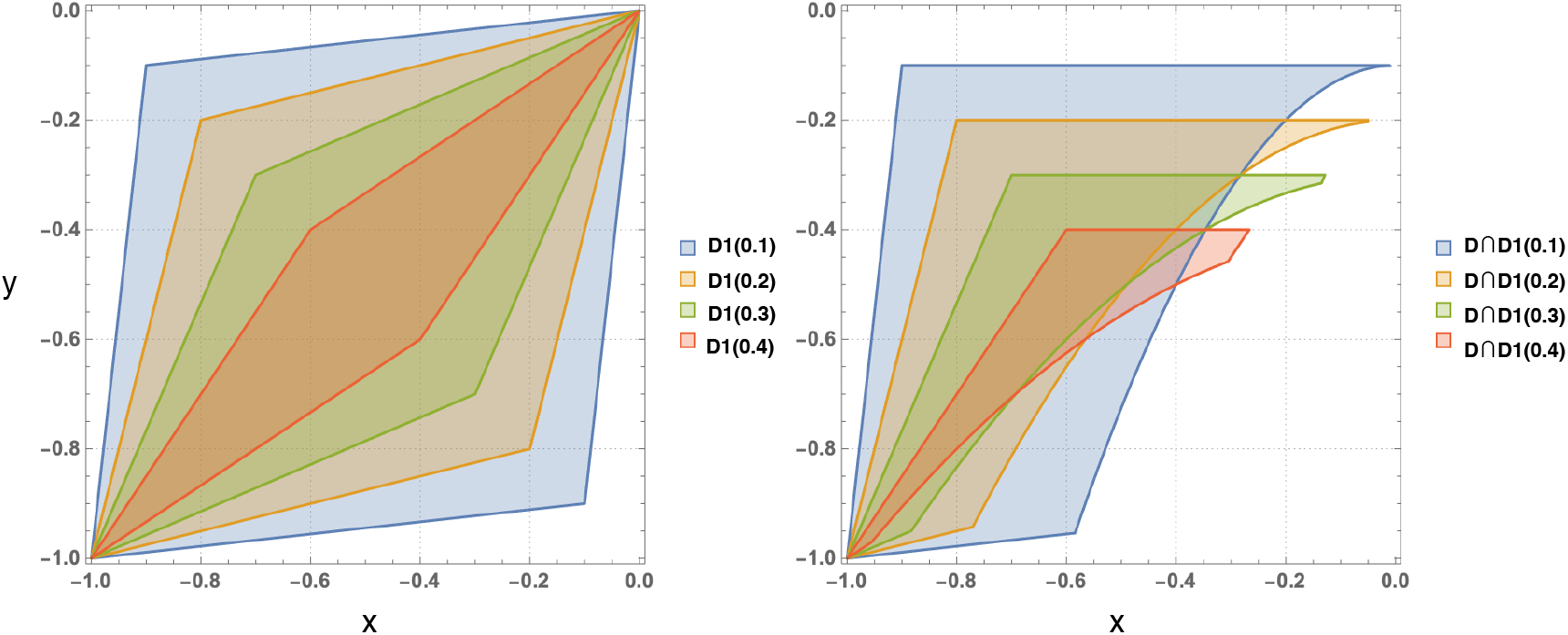
SH game: Regions *D*_1_ and *D* ∩ *D*_1_ are shown for different values of *q*, namely, *q* = 0.1, 0.2, 0.3 and 0.4.

According to (10), the joint probability distribution of (*X, Y*) is

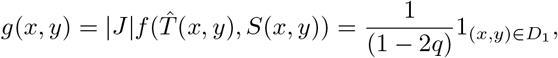

where

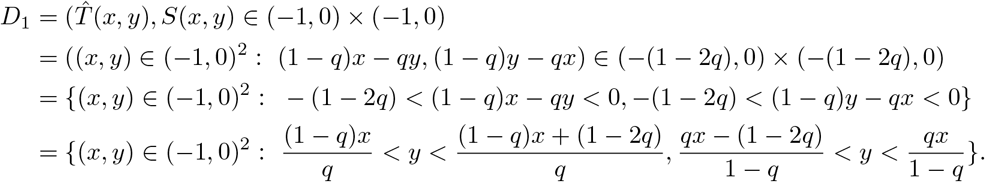

We can characterise *D*_1_ further by explicitly ordering the lower and upper bounds in the above formula. We have

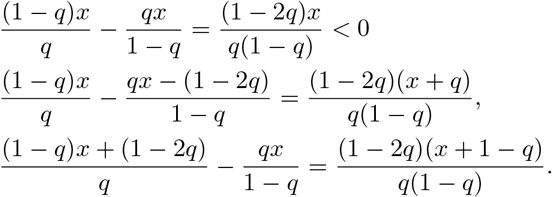

It follows that:

i. for −1 < *x* < −(1 − *q*) < −*q*:

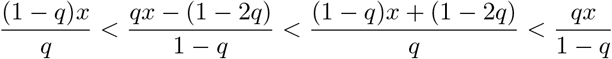
ii. For −(1 − *q*) < *x* < −*q*

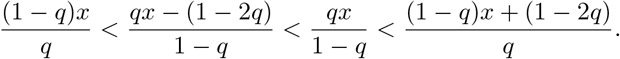
iii. for −*q* < *x* < 0

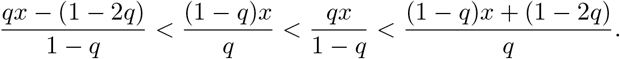

Hence

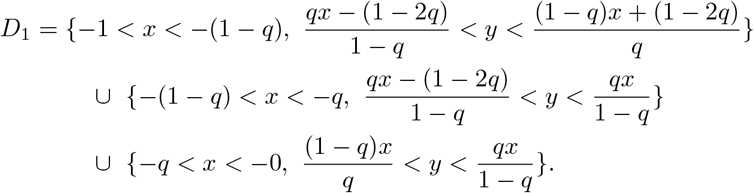

Thus, *D*_1_ is the quadrilateral *ABOC* with vertices

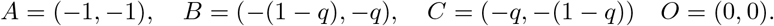

#### Theorem 2.3

*The probability that SH games have* 3 *equilibria is given by*

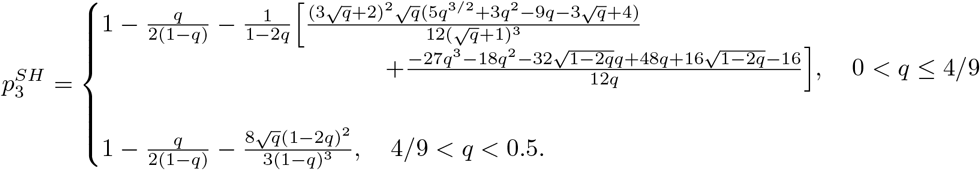

*Proof.* According to Proposition 2.2, in order to compute 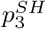 we need to compute the area of *D* ∩ *D*_1_. To this end we need to understand the intersections of the parabola 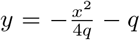 with the lines 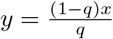 and 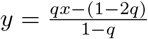. The parabola 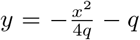 always intersects with the line 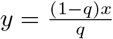 inside the domain (−1, 0)^2^ at the point

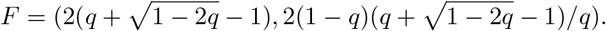

By comparing 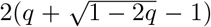 with −*q*, it follows that the point *F* is inside the edge *OC* if 0 ≤ *q* ≤ 4/9 and is outside *OC* whenever 4/9 < *q* < 0.5. On the other hand, the parabola 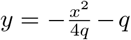 meets the line 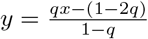 at two points 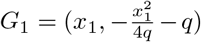 and 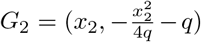 with

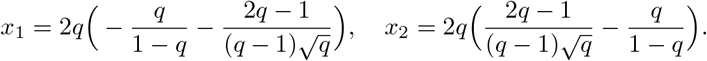

By comparing *x*_1_ and *x*_2_ with −*q* we have *G*_1_ is always in the edge *AC*, while *G*_2_ is outside *AC* if 0 < *q* ≤ 4/9 and is inside *AC* if 4/9 < *q* < 0.5.

In conclusion

1. **for** 0 < *q* ≤ 4/9: the intersection *D* ∩ *D*_1_ is the domain formed by vertices *A*, *B*, *E*, *F*, and *G*_1_ where 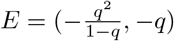 (which is the intersection of *y* = −*q* with 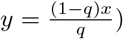 and

- *A* and *B* are connected by the line 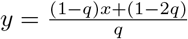,
- *B* and *E* are connected by the line *y* = −*q*,
- *E* and *F* are connected by the line 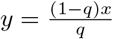,
- *F* and *G*_1_ are connected by the parabola 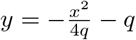,
- *G*_1_ and *A* are connected by the line 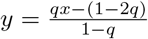.
2. **for** 4/9 < *q* < 0.5: the intersection *D* ∩ *D*_1_ is the domain formed by the vertices *A, B, E, C, G*_1_, *G*_2_ where

- *A* and *B* are connected by the line 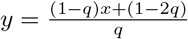,
- *B* and *E* are connected by the line y = −q,
- *E* and *C* are connected by the line 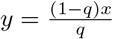
- *C* and *G*_1_ are connected by the line 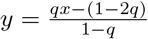,
- *G*_1_ and *G*_2_ are connected by the parabola 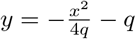,
- *G*_2_ and *A* are connected by the line 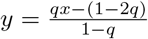.

See Figure 4 for illustrations of the two cases above, with *q* = 0.3 and *q* = 0.45 respectively. We are now ready to compute the probability that the SH game has three equilibria.

**Figure 4:**
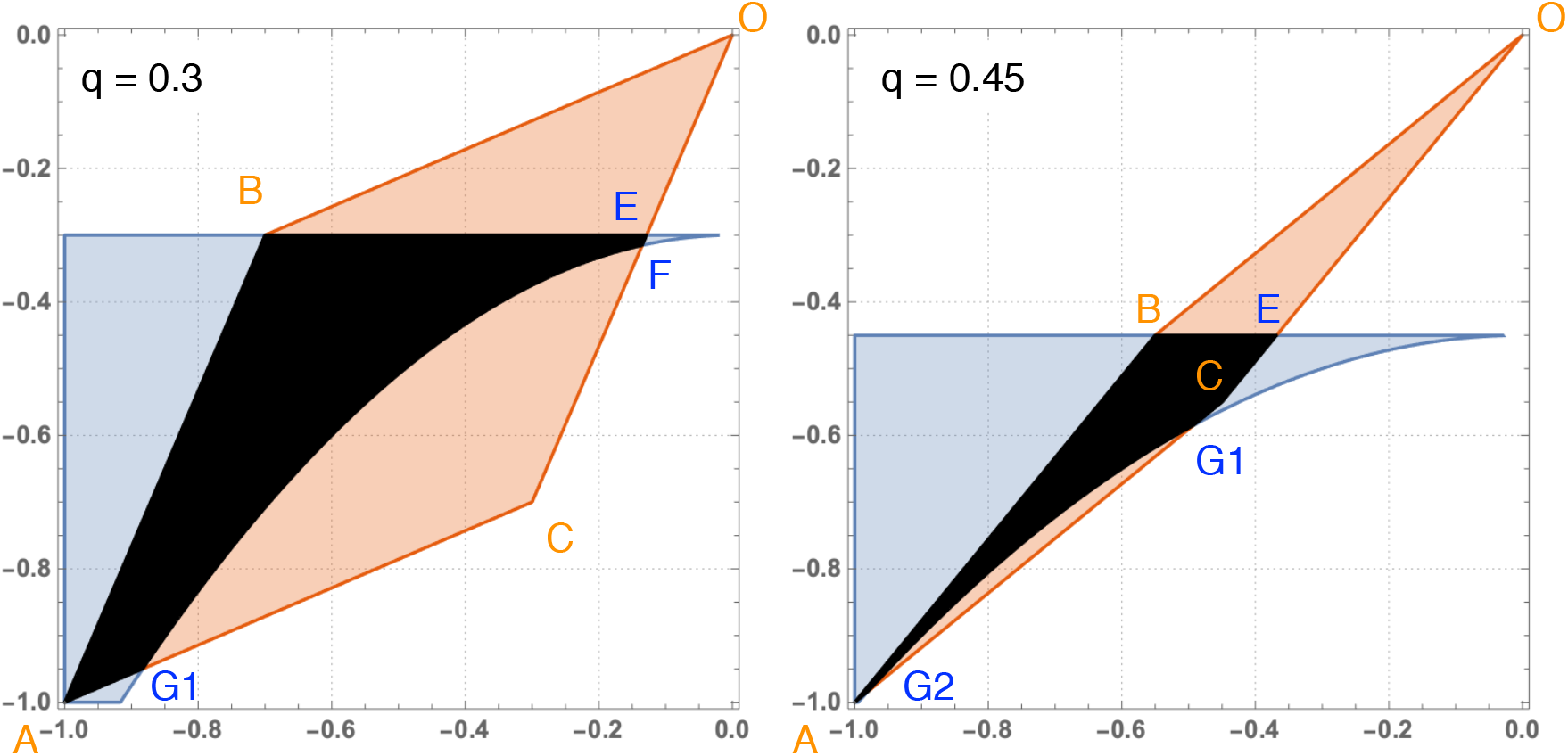
Details of *D* ∩ *D*_1_ for *q* = 0.3 and *q* = 0.45

**For** 0 < *q* ≤ 4/9: The probability that the SH game has three equilibria is

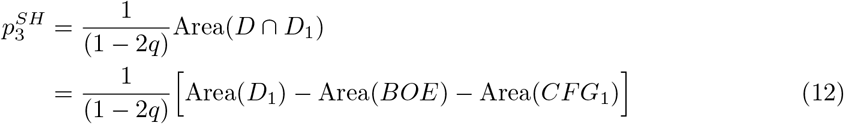

We proceed by computing the areas in the expression above. Area of *D*_1_ is

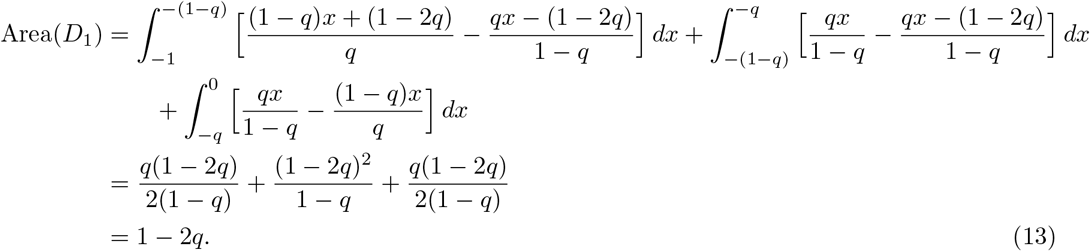

Area of *BOE* is

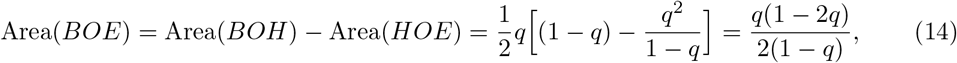

where *H* = (0, −*q*). Area of *CFG* is

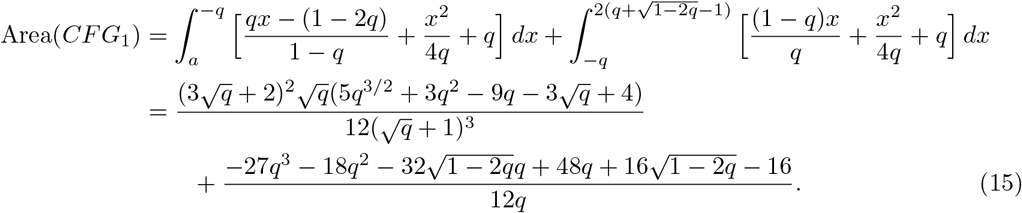

Substituting (13), (14), and (15) back to (12) we obtain, for 0 < *q* ≤ 4/9

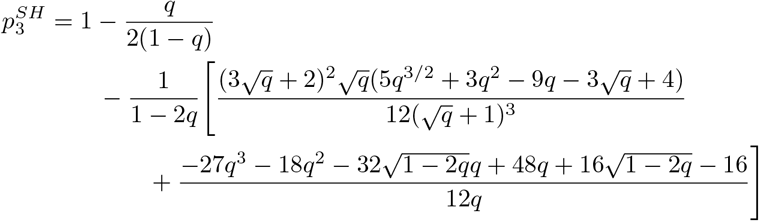

Now we consider the remaining case 4/9 < *q* < 0.5. In this case

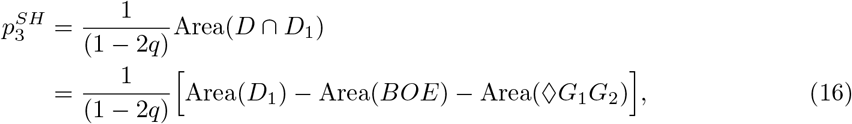

where ◊*G*_1_*G*_2_ is the domain with vertices *G*_1_ and *G*_2_ formed by the parabola 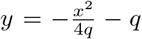 and the line 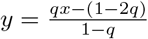. Thus

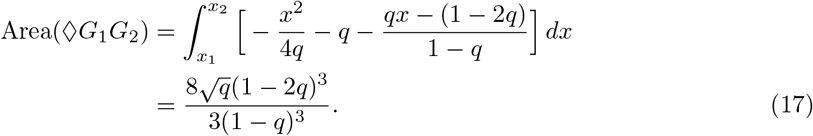

Substituting (13), (14) and (17) back to (16) we obtain, for 4/9 < *q* < 0.5,

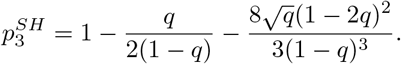

This finishes the proof of this theorem

### 2.4 Prisoner’s Dilemma (PD)

#### Proposition 2.4.

*The probability that PD games have* 3 *equilibria is given by*

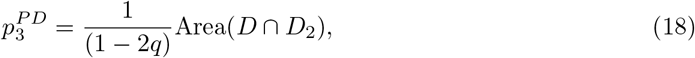

*where D is defined above (as in the case of the SH games) and D*_2_ *is the triangle MNO with vertices*

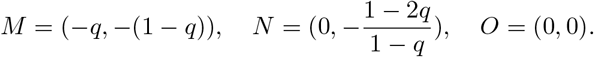

See Figure 5 for illustration of *D*_2_ and its intersection with *D* for several values of *q*.

**Figure 5:**
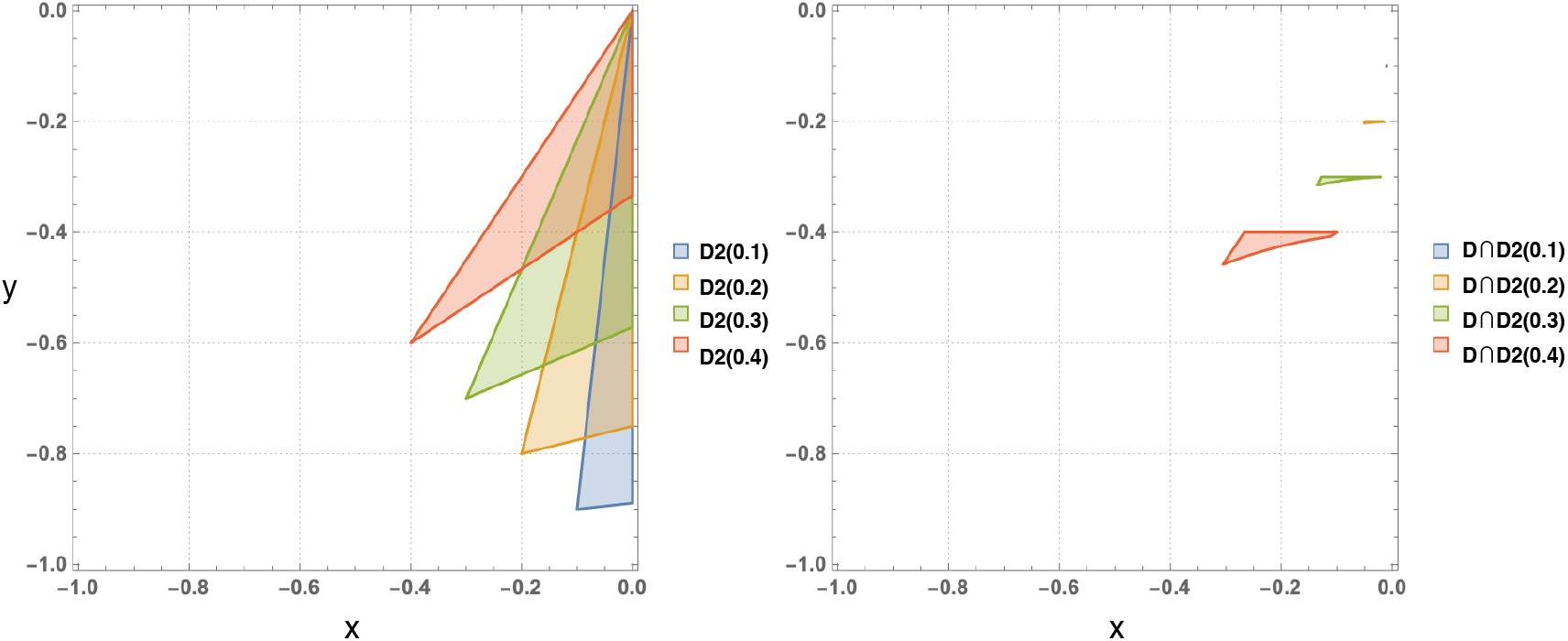
PD game: Regions *D*_2_ and *D* ∩ *D*_2_ are shown for different values of *q*, namely, *q* = 0.1, 0.2, 0.3 and 0.4.

*Proof.* Recall that in PD games we have *T* ~ *U* (1, 2), *S* ~ (−1, 0). Thus 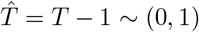. The joint distribution of 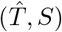 is

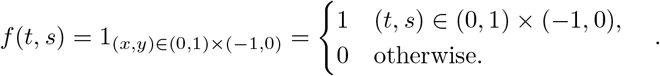

According to (10), the joint probability distribution of (*X, Y*) is

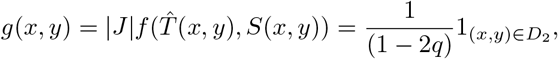

where

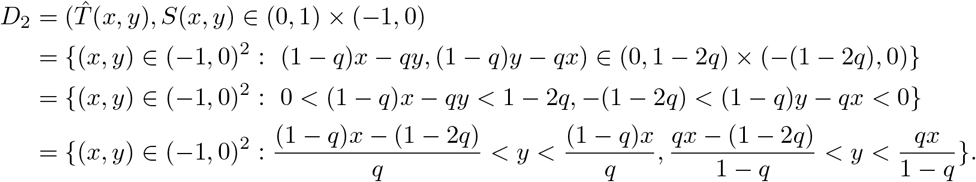

We now characterise *D*_2_ further. We have

1. for −1 < *x* < −*q* then

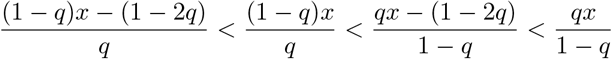
2. for −*q* < *x* < 0 then

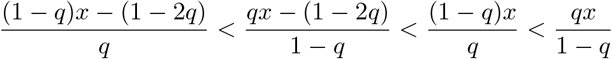

It follows that

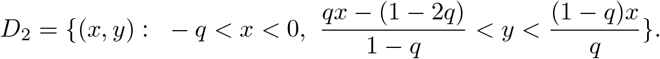

Thus *D*_2_ is the triangle *MNO* with vertices

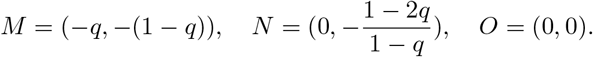

The probability that the SH game has three equilibria is thus

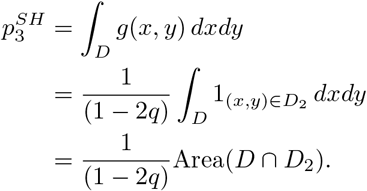

This finishes the proof of this proposition.

The following elementary lemma provides an upper bound for 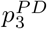, particularly implying that it tends to 0 as *q* goes to 0.

#### Lemma 2.5.

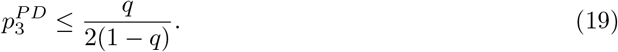

*As a consequence, 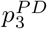 is always smaller or equal to* 0.5 *and tends to* 0 *as q tends to 0
Proof.* Area of *D*

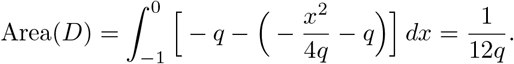

Area of *D*_2_

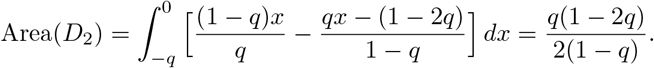

Hence

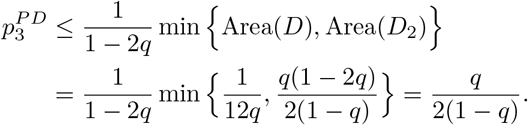

#### Theorem 2.6.

*The probability that PD games has three equilibria is given by*

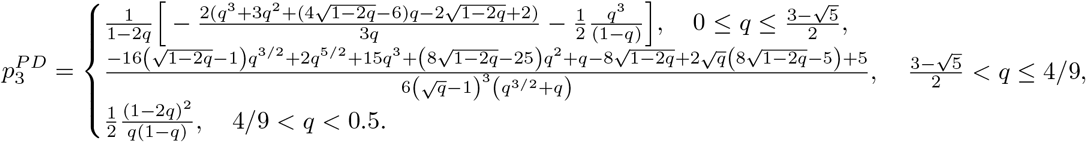

*Proof.* According to Proposition 2.4, in order to compute 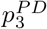 we need to compute the area of *D* ∩ *D*_2_. As in proof of Theorem 2.3, the parabola 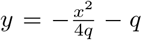 always intersects with the line 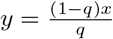 inside the domain (−1, 0)^2^ at the point

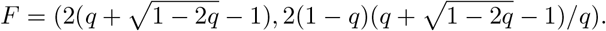

The point *F* is inside the edge *DC* if 0 ≤ *q* ≤ 4/9 and is outside *DC* whenever 4/9 < *q* < 0.5.

On the other hand, the parabola 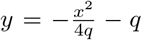 meets the line 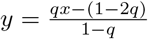 at two points 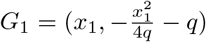 and 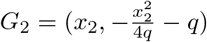 with

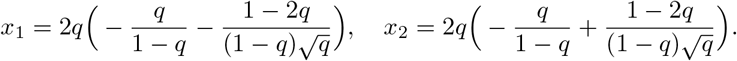

*G*_1_ is always outside edge *AC*. *G*_2_ is inside it if 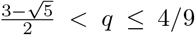 and outside it otherwise. Therefore, we have three cases (see Figure 6 for illustration).

**Figure 6:**
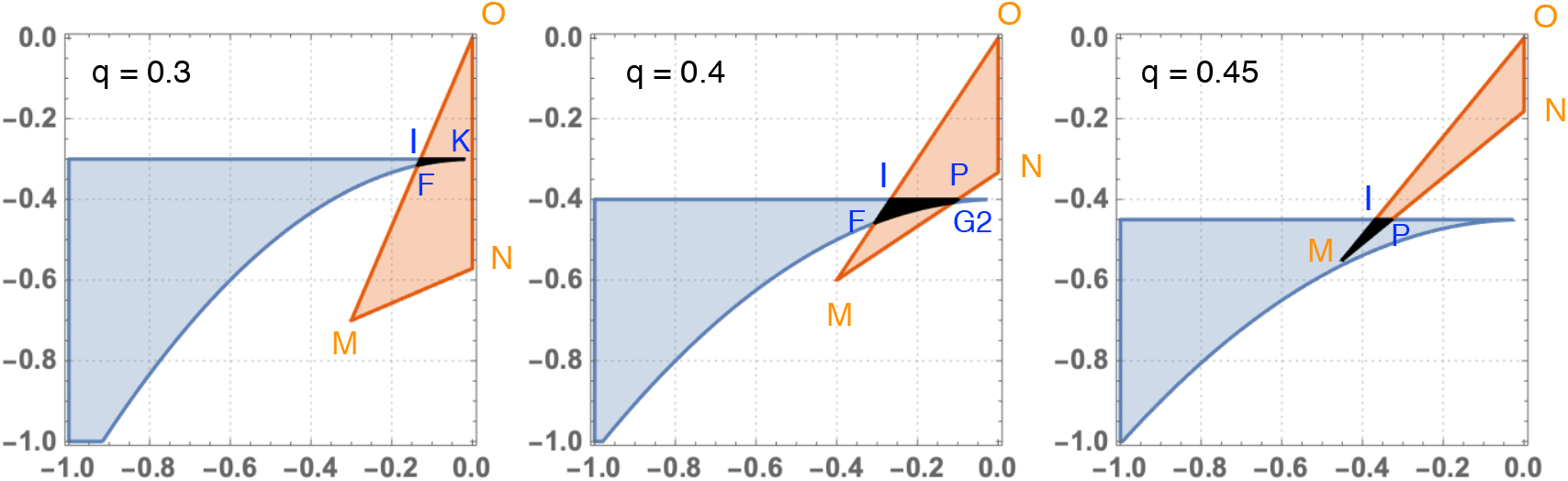
*D* ∩ *D*_2_ for *q* = 0.3, 0.4, 0.45

1. **For** 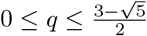: the intersection *D* ∩ *D*_2_ is formed by *I, F, K* where *I* = (−*q*^2^/(1 − *q*), −*q*) (which is the intersection of *y* = −*q* with 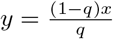), *K* = (0. − *q*) and

- *I* and *F* are connected by the line 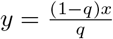,
- *F* and *K* are connected by the parabola 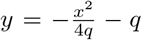,
- *K* and *I* are connected by the line *y* = −*q*. In this case

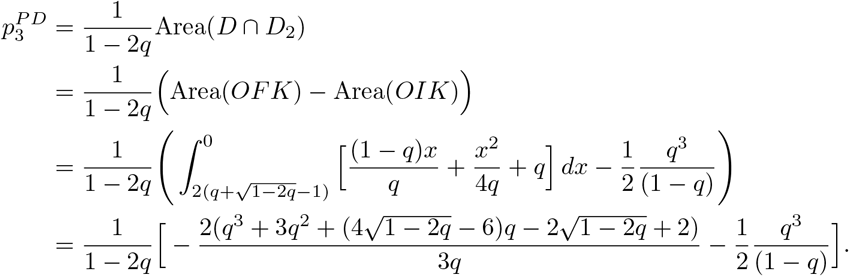
2. **For** 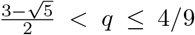: the intersection *D* ∩ *D*_2_ is formed by *F*, *I*, *P* = ((*q*^3^ − 3*q* + 1)/q, −*q*) (which is the intersection of y = −q with 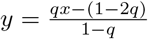 and *G*_2_, where

- *F* and *I* are connected by the line 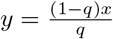,
- *I* and *P* are connected by the line y = −q,
- *P* and *G*_2_ are connected by the line 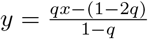,
- *G*_2_ and *F* are connected by the parabola 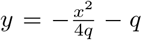. In this case

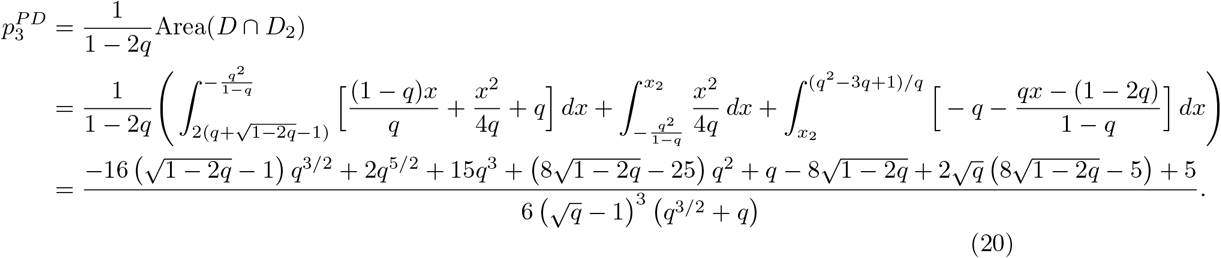
3. **For** 4/9 < *q* < 0.5: the intersection *D* ∩ *D*_2_ is the triangle *MIP* where *M* = (−*q*, −(1 − *q*))

- *M* and *I* are connected by the line 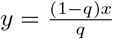,
- *I* and *P* are connected by the line *y* = −*q*,
- *P* and *M* are connected by the line 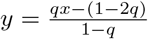. In this case

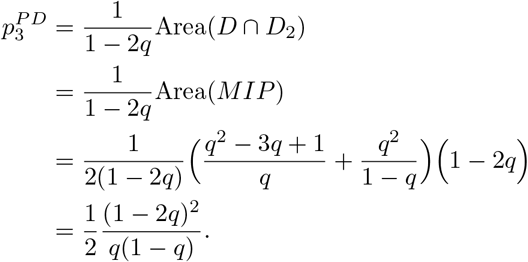

In conclusion

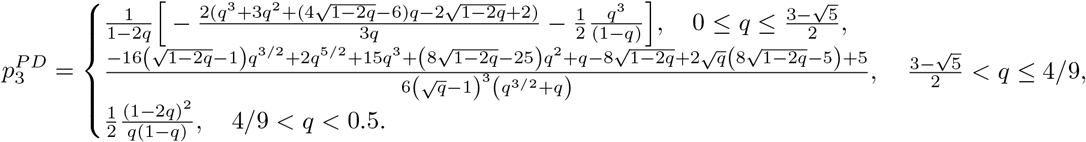

This completes the proof of this theorem.

## 3 Summary and outlook

It has been shown that in human behaviours and other biological settings, mutation is non-negligible [31, 25, 33]. How mutation affects the complexity and bio-diversity of the evolutionary systems is a fundamental question in evolutionary dynamics [20, 27]. In this paper, we have addressed this question for random social dilemmas by computing explicitly the probability distributions of the number of equilibria in term of the mutation probability. Our analysis based on random games is highly relevant and practical, because it is often the case that one might know the nature of a game at hand (e.g., a coordination or cooperation dilemma), but it is very difficult and/or costly to measure the exact values of the game’s payoff matrix. Our results have clearly shown the influence of the mutation on the number of equilibria in SH-games and PD-games. The probability distributions in these games are much more complicated than in SD-games and H-games and significantly depend on the mutation strength. For a summary of our results, see again Box 1 and Figure 1. Our analysis has made use of suitable changes of variables, which expressed the probability densities in terms of area of certain domains. For future work, we plan to generalise our method to other approaches to studying random social dilemmas such as finite population dynamics and payoff disturbances [13, 30, 1, 2], as well as to multi-player social dilemma games [23, 28, 18].

## Acknowledgments

TAH is also supported by Leverhulme Research Fellowship (RF-2020-603/9).

## References

[1] M. A. Amaral and M. A. Javarone. Heterogeneity in evolutionary games: an analysis of the risk perception. Proceedings of the Royal Society A: Mathematical, Physical and Engineering Sciences, 476(2237):20200116, 2020.

[2] M. A. Amaral and M. A. Javarone. Strategy equilibrium in dilemma games with off-diagonal payoff perturbations. Phys. Rev. E, 101:062309, Jun 2020.

[3] M. Broom. Evolutionary games with variable payoffs. C. R. Biologies, 328, 2005.

[4] M. Broom, C. Cannings, and G. T. Vickers. On the number of local maxima of a constrained quadratic form. Proc. R. Soc. Lond. A, 443:573–584, 1993.

[5] M. H. Duong and T. A. Han. On the expected number of equilibria in a multi-player multistrategy evolutionary game. Dynamic Games and Applications, pages 1–23, 2015.

[6] M. H. Duong and T. A. Han. Analysis of the expected density of internal equilibria in random evolutionary multi-player multi-strategy games. Journal of Mathematical Biology, 73(6):1727–1760, 2016.

[7] M. H. Duong and T. A. Han. On equilibrium properties of the replicator–mutator equation in deterministic and random games. Dynamic Games and Applications, 10(3):641–663, Sep 2020.

[8] M. H. Duong, H. M. Tran, and T. A. Han. On the distribution of the number of internal equilibria in random evolutionary games. Journal of Mathematical Biology, 78(1):331–371, Jan 2019.

[9] R. Durrett. The Essentials of Probability. Duxbury Press, 1994.

[10] C. S. Gokhale and A. Traulsen. Evolutionary games in the multiverse. Proc. Natl. Acad. Sci. U.S.A., 107(12):5500–5504, 2010.

[11] K. P. Hadeler. Stable polymorphisms in a selection model with mutation. SIAM Journal on Applied Mathematics, 41(1):1–7, 1981.

[12] T. A. Han, A. Traulsen, and C. S. Gokhale. On equilibrium properties of evolutionary multiplayer games with random payoff matrices. Theoretical Population Biology, 81(4):264 – 272, 2012.

[13] W. Huang and A. Traulsen. Fixation probabilities of random mutants under frequency dependent selection. J. Theor. Biol., 263:262–268, 2010.

[14] L. A. Imhof, D. Fudenberg, and M. A. Nowak. Evolutionary cycles of cooperation and defection. Proceedings of the National Academy of Sciences, 102(31):10797–10800, 2005.

[15] N. L. Komarova. Replicator–mutator equation, universality property and population dynamics of learning. Journal of Theoretical Biology, 230(2):227 – 239, 2004.

[16] N. L. Komarova and S. A. Levin. Eavesdropping and language dynamics. Journal of Theoretical Biology, 264(1):104 – 118, 2010.

[17] N. L. Komarova, P. Niyogi, and M. A. Nowak. The evolutionary dynamics of grammar acquisition. Journal of Theoretical Biology, 209(1):43 – 59, 2001.

[18] Q. Luo, L. Liu, and X. Chen. Evolutionary dynamics of cooperation in the n-person stag hunt game. Physica D: Nonlinear Phenomena, 424:132943, 2021.

[19] R. M. May. Stability and complexity in model ecosystems, volume 6. Princeton university press, 2001.

[20] M. A. Nowak. Evolutionary Dynamics. Harvard University Press, Cambridge, MA, 2006.

[21] M. A. Nowak, N. L. Komarova, and P. Niyogi. Evolution of universal grammar. Science, 291(5501):114–118, 2001.

[22] R. Olfati-Saber. Evolutionary dynamics of behavior in social networks. In 2007 46th IEEE Conference on Decision and Control, pages 4051–4056, Dec 2007.

[23] J. M. Pacheco, F. C. Santos, M. O. Souza, and B. Skyrms. Evolutionary dynamics of collective action in n-person stag hunt dilemmas. Proceedings of the Royal Society of London B: Biological Sciences, 276(1655):315–321, 2009.

[24] D. Pais, C. Caicedo-Núnẽz, and N. Leonard. Hopf bifurcations and limit cycles in evolutionary network dynamics. SIAM Journal on Applied Dynamical Systems, 11(4):1754–1784, 2012.

[25] David G Rand, Corina E Tarnita, Hisashi Ohtsuki, and Martin A Nowak. Evolution of fairness in the one-shot anonymous ultimatum game. Proceedings of the National Academy of Sciences, 110(7):2581–2586, 2013.

[26] F. C. Santos, J. M. Pacheco, and T. Lenaerts. Evolutionary dynamics of social dilemmas in structured heterogeneous populations. Proc. Natl. Acad. Sci. U.S.A., 103:3490–3494, 2006.

[27] K. Sigmund. The calculus of selfishness. Princeton Univ. Press, 2010.

[28] M. O. Souza, J. M. Pacheco, and F. C. Santos. Evolution of cooperation under n-person snowdrift games. Journal of Theoretical Biology, 260(4):581 – 588, 2009.

[29] P. F. Stadler and P. Schuster. Mutation in autocatalytic reaction networks. Journal of Mathematical Biology, 30(6):597–632, Jun 1992.

[30] A. Szolnoki and M. Perc. Seasonal payoff variations and the evolution of cooperation in social dilemmas. Scientific reports, 9(1):1–9, 2019.

[31] A. Traulsen, C. Hauert, H. De Silva, M. A Nowak, and K. Sigmund. Exploration dynamics in evolutionary games. Proc. Natl. Acad. Sci. USA, 106:709–712, 2009.

[32] Z. Wang, S. Kokubo, M. Jusup, and J. Tanimoto. Universal scaling for the dilemma strength in evolutionary games. Physics of life reviews, 14:1–30, 2015.

[33] I. Zisis, S. Di Guida, T. A. Han, G. Kirchsteiger, and T. Lenaerts. Generosity motivated by acceptance-evolutionary analysis of an anticipation game. Scientific reports, 5(1):1–11, 2015.

